## supplemental figures for "Top-down acetylcholine contributes to social discrimination via enabling action potentials in olfactory bulb vasopressin cells"

### Supplemental information

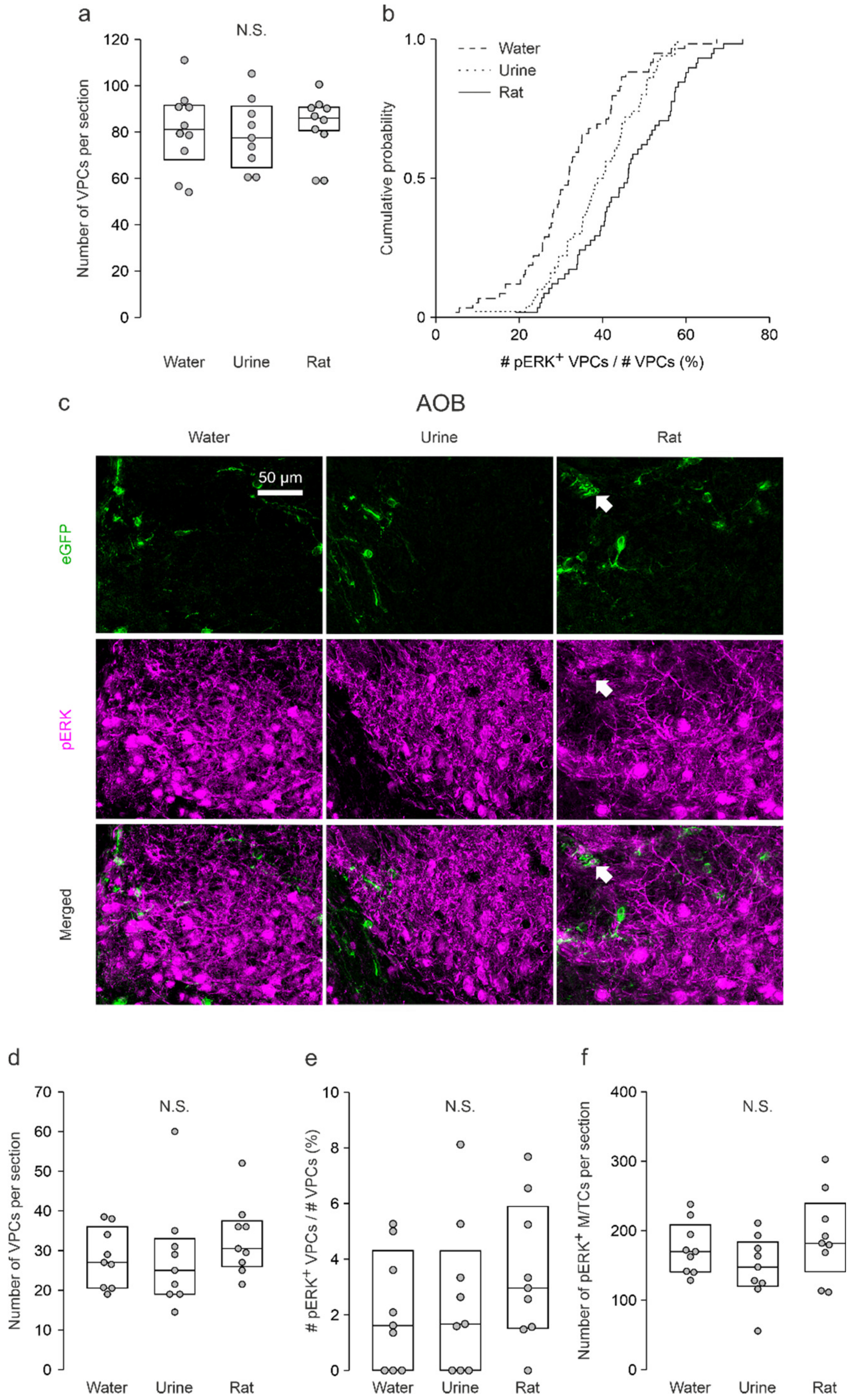


Supplemental Fig. 1 VPC activation in the MOB and AOB
(a) Averaged number of MOB VPCs per section in different stimulation groups. (b) Cumulative probability of fraction of number of pERK$^+$ MOB VPCs over number of MOB VPCs in each sections not averaged over animals from different stimulation groups. 59 sections (water), 50 sections (urine), 58 sections (rat). (c) Representative average z-projections of the accessory olfactory bulb that were immune-stained for eGFP (green, CF488) and pERK (magenta, CF 594). Arrows indicate a cell that are double-labeled for eGFP and pERK. Scale bar, 50 µm. (d+f) Averaged number of AOB VPCs or of pERK$^+$ AOB M/TCs per section in different stimulation groups. (e) Averaged fraction of number of pERK$^+$ AOB VPCs over number of AOB VPCs in different stimulation groups (%). Data are presented as box-plots including median and distribution of single data points. ANOVA, N.S., not significant. n=9 (water), n=9 (urine), n=9 (rat).

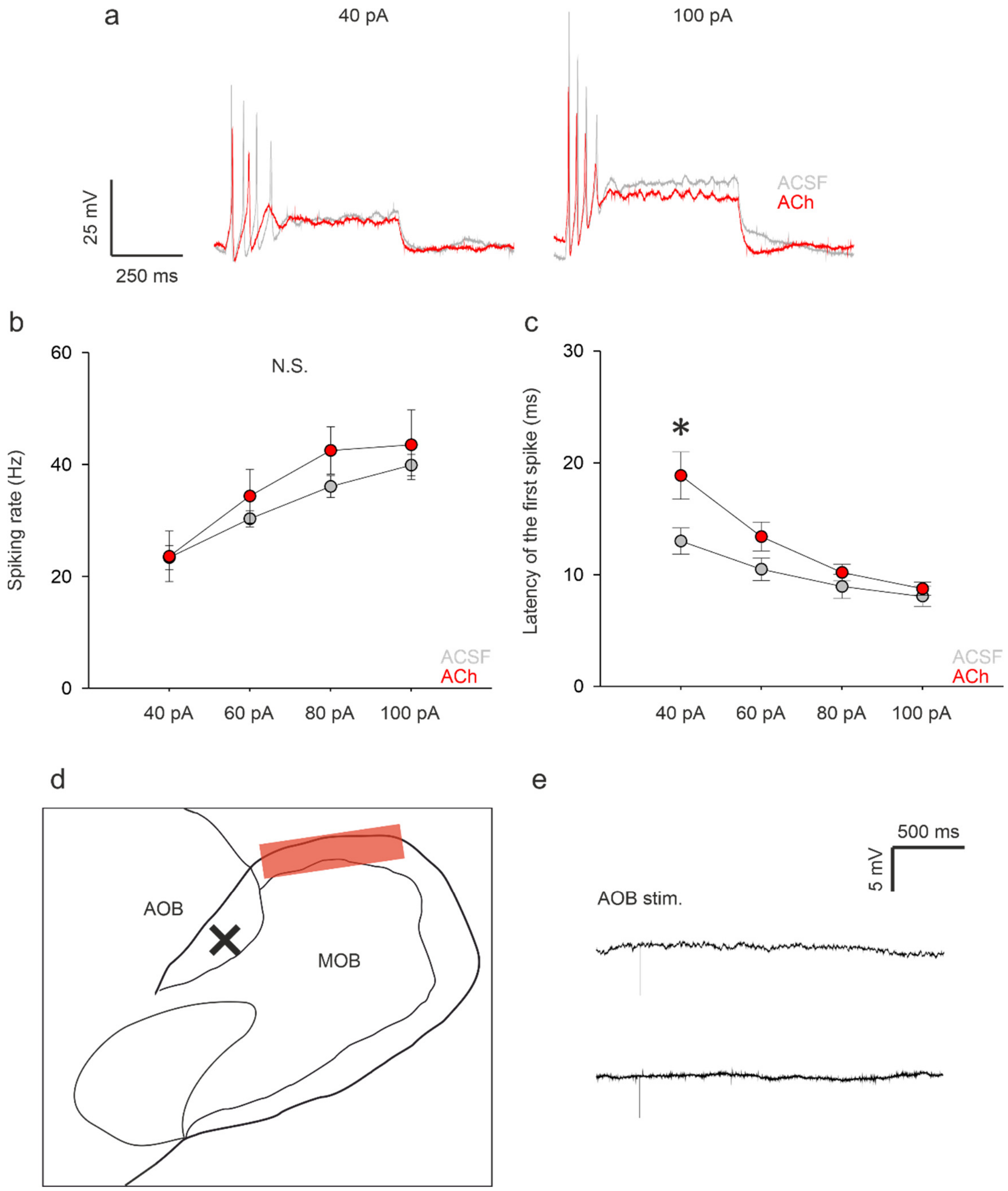


Supplemental Fig. 2 **ACh does not alter intrinsic excitability in VPCs and electrical AOB stimulation does not evoke excitatory responses in MOB VPCs**

(a) Representative traces of responses to somatically applied current steps in the ACSF condition (grey) and during bath application of ACh (100 µM, red). (b+c) Spiking rates of action potential trains (left) and latency of the first spike (right) evoked by somatic current injection (40-100 pA) in the ACSF condition (grey) and during bath application of ACh (100 µM, red). (2) × (2) mixed model ANOVA (intensity [within subject] × treatment [within-subject]). N.S., not significant. LSD for single comparison, *$p<0.05$ ACh (40pA) vs. ACSF (40pA). Data are means ± SEM. (d) Schematic drawing of the sagittal OB. The cross indicates where the stimulation electrode was positioned. The red bar indicates the dorsal region of the MOB where patch-clamp recordings from MOB VPCs were performed. (e) Representative averaged trace of response from MOB VPCs to electrical vomeronasal nerve/ AOB stimulation (50-500 µA, 100 µs).

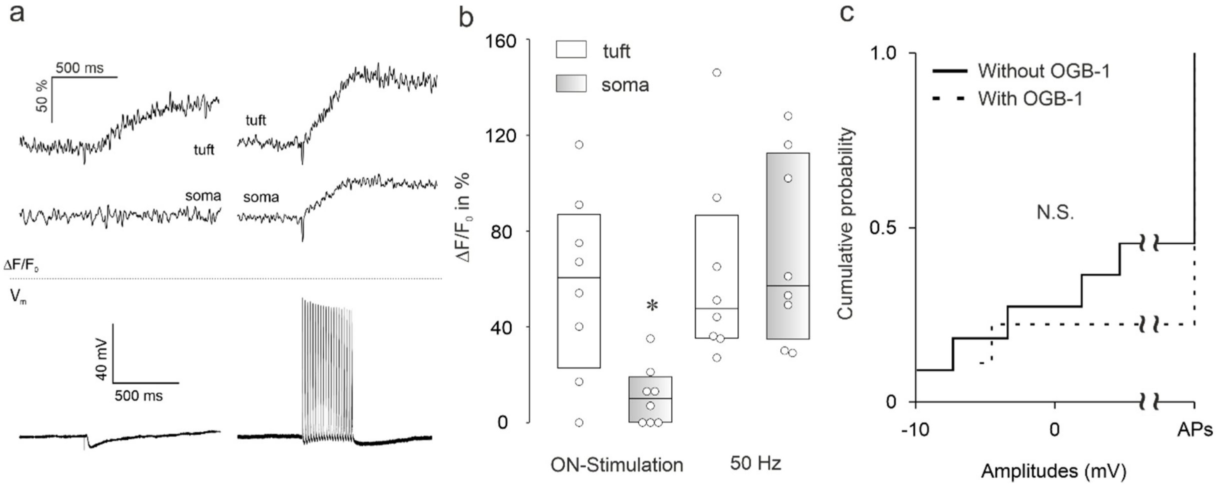


Supplemental Fig. 3 **Increase of intracellular Ca2+ in the tuft, but not near the soma following olfactory nerve stimulation** (a) Representative averaged (n=3) ΔF/F transients in response to a single olfactory nerve stimulation (ON, 50-100 µA, 100 µs) and somatically evoked 50 Hz trains (20 APs, 50 Hz, 400 ms). (b) Cumulative representation of ΔF/F following ON or 50Hz train stimulation measured at the tuft and at the apical dendrite near the soma. Data are presented as box-plots including median and distribution of single data points. * p = 0.002 vs. tuft; Paired t-test. (c) Cumulative probability of evoked PSP amplitudes in the ACh condition with or without OGB-1 in the intracellular solution (n=9/11). The amplitudes of APs were set as 100 mV. Kruskal-Wallis test for variation comparison.

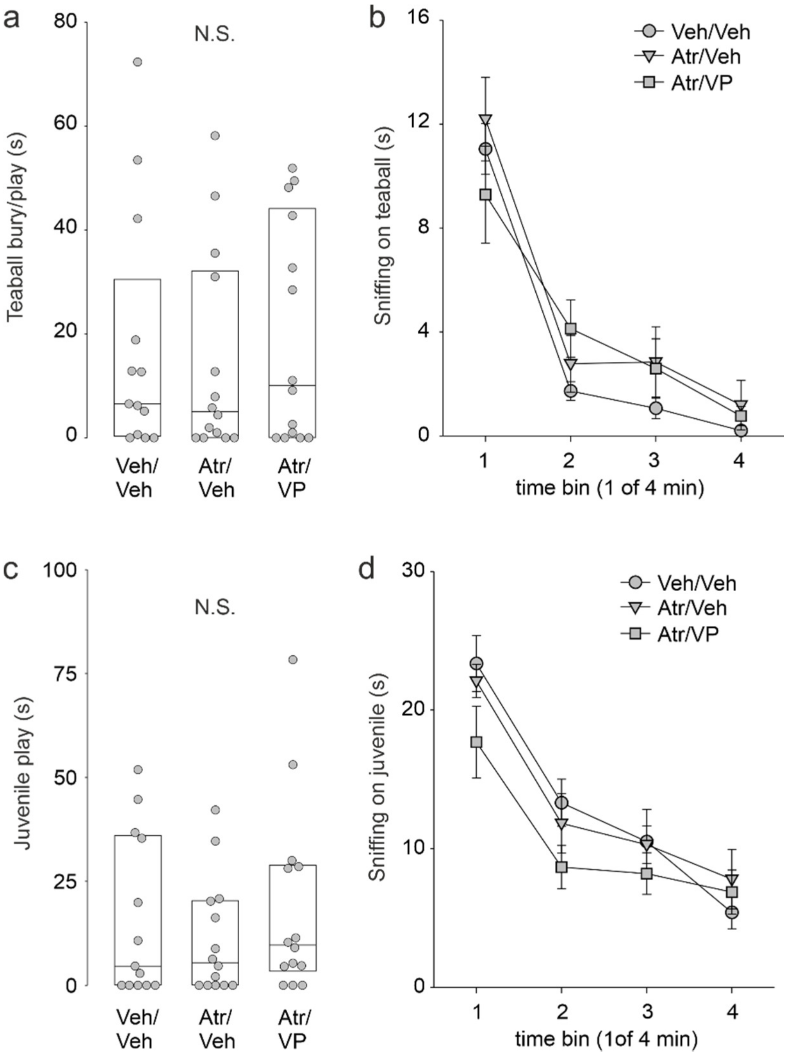


Supplemental Fig. 4 **Atropine and VP microinjection into the OB does not interfere with non-social investigatory/play behavior and habituation.**

(a+c) Amount of time (s) that rats are engaged in burying/playing with the teaball during neutral odor stimulation (amylacetate or carvone) and playing with the juvenile stimulus rat during the sampling phase of the social discrimination test. Data are presented as box-plots including median and distribution of single data points. Kruskal-Wallis Test, n=13 (Veh/Veh), n=14 (Atr/Veh, 1µg), n=14 (Atr/VP, 1µg/1ng)$_{c1,e1}$. (b+d) Amount of time (s) within time bins of 1 min rats investigate the teaball during neutral odor presentation and the juvenile stimulus rat during the sampling phase of the social discrimination test. Data are presented as means ± SEM. (4) × (3) mixed model ANOVA (time bin [within subject] × treatment [within subject]), n=13 (Veh/Veh), n=14 (Atr/Veh, 1µg), n=14 (Atr/VP, 1 µg/1ng)$_{d1,f1}$.
